# Nicotinic receptors atlas in the adult human lung

**DOI:** 10.1101/2020.06.29.176750

**Authors:** Zania Diabasana, Jeanne-Marie Perotin, Randa Belgacemi, Julien Ancel, Pauline Mulette, Gonzague Delepine, Philippe Gosset, Uwe Maskos, Myriam Polette, Gaëtan Deslée, Valérian Dormoy

## Abstract

Nicotinic acetylcholine receptors (nAChRs) are pentameric ligand-gated ion channels responsible for the rapid neural and neuromuscular signal transmission. Although it is well documented that 16 subunits are encoded by the human genome, their presence in airway epithelial cells (AEC) remains poorly understood, and contribution to pathology is mainly discussed in the context of cancer. We analysed nAChR subunit expression in the human lung of smokers and non-smokers using transcriptomic data for whole lung tissues, isolated large AEC, and isolated small AEC. We identified differential expressions of nAChRs in terms of detection and repartition in the three modalities. Smoking-associated alterations were also unveiled. Then, we identified a nAChR transcriptomic print at a single cell level. Finally, we reported the localizations of detectable nAChRs in bronchi and large bronchioles. Thus, we compiled the first complete atlas of pulmonary nAChRs to open new avenues to further unravel the involvement of these receptors in lung homeostasis and respiratory diseases.

## INTRODUCTION

Nicotinic acetylcholine receptors (nAChRs) are ligand-gated (cation-permeable) proteins expressed in the brain and in non-neuronal cells including lung AEC, macrophages, neutrophils, and muscle cells [1]. These receptors are composed of five subunits organized as homo- or hetero-pentamers forming a channel permeable to cations (predominantly Na^+^, K^+^ and Ca^2+^) [1,2]. Each subunit shares a common structure comprising a large amino-terminal extracellular domain (about 200 residues), four transmembrane domains M1, M2, M3, and M4 (less than 30 residues), and a variable cytoplasmic domain (90 to 270 residues) [3–5]. There are 16 mammalian subunits, namely CHRNA1-A7, A9–A10, B1–B4, D, G and E [2]. Muscle-type nAChRs are generally assembled from 2(CHRNA1)/B1/G/D or 2(CHRNA1)/B1/E/D subtypes depending on muscle innervation [5–8]. Neuronal nAChRs are assembled from CHRNA2-A10 and CHRNB2-B4, and can either form homopentamers solely composed of CHRNA7 or CHRNA9 subunits, or heteropentamers constituted of CHRNA2-A6 and CHRNA10 combined with CHRNB2-B4 subunits [3,9–13]. All these subtypes confer different functional properties to the receptors [3,9,11].

In the lung, acetylcholine binding to nAChR leads to an allosteric conformational change permitting the channel opening, followed by ion fluxes that participate in cell survival, differentiation, and proliferation [14,15]. Nicotine, one of the components of cigarette smoke, acts as an agonist implicated in the inhibition of apoptosis and oxidative stress responses ultimately leading to lung impairments due to long-term exposure [5,16,17]. Previous studies have established that multiple single nucleotide nAChR polymorphisms are respectively associated with risks of lung cancer and chronic obstructive pulmonary disease (COPD), highlighting their potential implication in respiratory diseases [14,18,19]. In addition, it has been hypothesized that the nAChRs may play a role in coronavirus disease 2019 (COVID-19) and might represent a therapeutic target, particularly regarding their potential contribution in the regulation of angiotensin converting enzyme-2 (ACE-2), the main receptor for severe acute respiratory syndrome coronavirus (SARS-CoV-2) [20–22].

If the general expressions of muscle and neuronal nAChRs are well known, little information is available regarding their expression in the lung and particularly in different AEC [23]. It is of the utmost importance to provide an atlas of pulmonary nAChRs because of their potential implication in respiratory diseases. Therefore, we conducted a transcriptomic and a proteomic analysis of the localization and expression of all human nAChRs in the adult lung.

## METHODS

### Human subjects

Patients scheduled for lung resection for cancer (University Hospital of Reims, France) were prospectively recruited (n=10) following standards established and approved by the institutional review board of the University Hospital of Reims, France (IRB Reims-CHU 20110612). In addition, 10 patients who underwent a routine large airway fiberoptic bronchoscopy with bronchial brushings under local anaesthesia according to international guidelines were also recruited (5 non-smokers, 5 smokers) [24]. Informed consent was obtained from all the patients. Patients with chronic obstructive pulmonary disease, asthma, cystic fibrosis, bronchiectasis or pulmonary fibrosis were excluded. At inclusion, age, sex, smoking history, and pulmonary function tests results were recorded. Ex-smokers were considered for a withdrawal longer than 5 years.

### Sample processing

Fresh airway epithelial cells (AEC) obtained from bronchial brushings (right lower lobe) were suspended within 30 min in RPMI (1% penicillin/streptomycin+ 10% BSA) before centrifugation (12,500 rpm x2 times). Cell pellet was dissociated in 1mL of Trypsine Versène^®^, centrifuged (12,500 rpm x2 times) and kept at -20°C until further steps.

### RT-qPCR analyses

Total RNA from AEC bronchial brushings was isolated by High Pure RNA isolation kit (Roche Diagnostics) and 250ng was reverse transcribed into cDNA by Transcriptor First Stand cDNA Synthesis kit (Roche Diagnostics). Quantitative PCR reactions were performed with fast Start Universal Probe Master kit and UPL-probe system in a LightCycler 480 Instrument (Roche Diagnostics) as recommended by the manufacturer. Primers used are listed in ***Supplementary Table S1***. Results for all expression data regarding transcripts were normalized to the expression of the house-keeping gene GAPDH amplified with the following primers: forward 5′-ACCAGGTGGTCTCCTCTGAC-3′, reverse 5′-TGCTGTAGCCAAATTCGTTG-3′. We verified that GAPDH transcript detection levels were highly similar between non-smokers and smokers to validate the housekeeping gene (average Ct=25.54±0.17 in non-smokers vs 25.35±0.34 in smokers; p=0.64). Relative gene expression was assessed by the ΔΔCt method [25] and expressed as 2^-ΔΔCt^. To compare data generated via PCR with RNAseq analysis, we transformed the transcript expressions in a percentage scale considering the highest and lowest values per subunit for the detection or across all the subunits for the repartition.

### Immunofluorescent staining and analyses

Immunohistochemistry was performed on formalin-fixed paraffin-embedded (FFPE) lung tissues distant from the tumour as previously described [26]. Only patients having no respiratory diseases were included (smokers and ex-smokers). Five μm sections were processed for H&E staining and observed on microscope (x20) to confirm the presence of bronchi and large bronchioles (pseudostratified epithelia). Bronchial epithelium was analysed on the entire slide including 2 to 7 units per patient. FFPE lung tissues sections slides were deparaffinised and blocked with 10% BSA in PBS for 30 min at room temperature. Tissue sections were then incubated with the primary antibodies as listed in ***Supplementary Table S2*** for one night at 4°C in 3% BSA in PBS. After PBS wash, a second primary antibody was used to highlight non-differentiated cells on epithelia for 2h at room temperature: p63 (AF1916, R&D Systems) or pan-cytokeratin (CK, E-AB-33599, Elabscience). Sections were washed with PBS and incubated with the appropriate secondary antibodies in 3% BSA in PBS for 30 min at room temperature. DNA was stained with DAPI during incubation with the secondary antibodies. Micrographs were acquired on a Zeiss AxioImageur (20x Ph) with ZEN software (8.1, 2012) and processed with ImageJ (National Institutes of Health) for analysis. For each patient, five random fields per section containing bronchi and large bronchioles were taken to evaluate the localization of nicotinic receptors on epithelial and stromal cells. We selected the most suitable primary antibodies directed against each subunit, considering external validations, identity and staining optimization to highlight the localization of nAChRs on bronchi and large bronchioles.

### RNAseq data analysis

Gene expression of non-smoking and smoking subjects with no chronic respiratory diseases were collected from datasets available online (GEO database; http://www.ncbi.nlm.nih.gov/geo) including whole lung tissues samples in 153 subjects (42 non-smokers, 111 smokers; GSE103174, 76925 and 47460) or small airway bronchoscopic samples in 135 subjects (63 non-smokers, 72 smokers; GSE11784).

In order to compare transcriptomic data extracted from various datasets or PCR reactions, we formatted the absolute values to a percentage scale. Concerning the detection of genes, we first identified for each gene the highest and the lowest expression values in both non-smokers and smokers to set the maximal value at 100%. After proportionally expressing each of the single expression value for all the subunits, the average was calculated and plotted in a graph. To discuss the relative level of expressions, we arbitrarily categorized 4 groups: (1) very high expressions, average percentage of expression is over 75% of the maximum; (2) high expressions, average percentage of expression is between 50 and 75% of the maximum; (3) moderate expressions, average percentage of expression is between 25 and 50% of the maximum; (4) and low expression, average percentage of expression is below 25% of the maximum.

Concerning the repartition, total expression of absolute values for all nAChR were summed for each patient of the considered data set to express the proportion of each subunit. The comparative average percentage of expression of each subunit for all patients was plotted in a pie chart.

### Single cell sequencing

The published dataset can be found on lungcellatlas.org and https://www.covid19cellatlas.org. We retained cell clustering based on the original studies and considered only lung samples from subjects with no respiratory disease [27]. An Illumina Hiseq 4000 per 10x Genomics chip position was used (n=6; 2,000-5,000 cells/sample). Additional sequencing was performed to obtain coverage of at least mean coverage of 100,000 reads per cell.

### Statistics

The data are expressed as mean values and percentages. Differences between groups were determined using the Student *t* test. A p-value <0.05 was considered significant.

## RESULTS

### Smoking-associated pulmonary nAChRs subunits transcripts expressions

Considering the whole lung containing all types of tissues (***Figure 1A***), the 16 nAChRs were detected among non-smoker subjects except for CHRNA7, that was consistent in both datasets containing non-smokers (***Figure 1B***). CHRNB1/E were very highly expressed; CHRNA6/A9/A10/B3/D were highly expressed. There was a significant increase of CHRNA1/A2/A7/B3/B4 transcript levels in smokers compared to non-smokers. Interestingly, CHRNA7 was only detected in smokers. On the contrary, CHRNA3/A4/A9/B2/D/G transcript levels were significantly decreased in smokers. The global repartition in non-smokers and smokers favoured CHRNA10/B1/E, representing almost half of all nAChRs expressed in lung tissues (***Figure 1C***). The differential repartitions of nAChRs between non-smokers and smokers matched their differential expressions.

**Figure 1.**
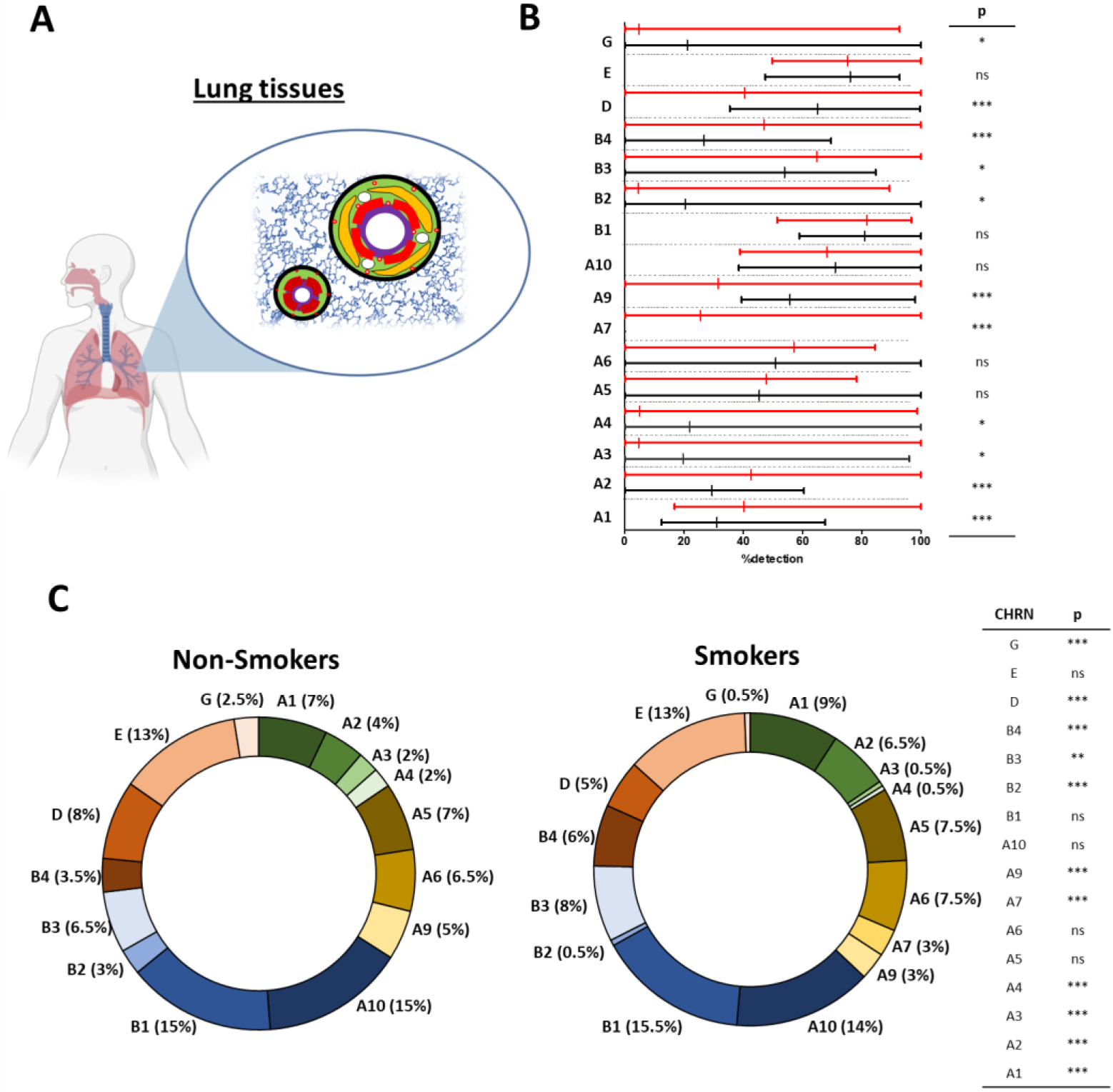
Evaluation of nAChR transcript levels in human lung tissues. **(A)**. Illustrations depicting the origin of the samples. Whole lung tissues contained all the tissues present in the lung either in parenchyma or in/around bronchi and bronchioles **(B)**. Histograms showing the detection of nAChRs in non-smokers (black) and smokers (red). **(C)**. Pie charts showing the repartition of nAChRs in non-smokers (left) and smokers (right). *p<0.05; **p<0.01; ***p<0.001 non-smokers (n=42) vs smokers (n=111).

In large AEC (LAEC), (***Figure 2A***) CHRNA1/A2/A4/B1/B3/B4/D were not detected in non-smokers (***Figure 2B***). CHRNA5 was very highly expressed; CHRNA7/A10 were highly expressed. Interestingly, CHRNB1/B4 were only detected in smokers. There was a significant decrease of CHRNA5/A10 transcript levels in smokers when compared to non-smokers. The global repartition in non-smokers and smokers favoured CHRNA5/A7/A9/A10 representing more than 75% of all nAChRs expressed in LAEC (***Figure 2C***). There were no significant differences in terms of nAChRs repartitions between non-smokers and smokers.

**Figure 2.**
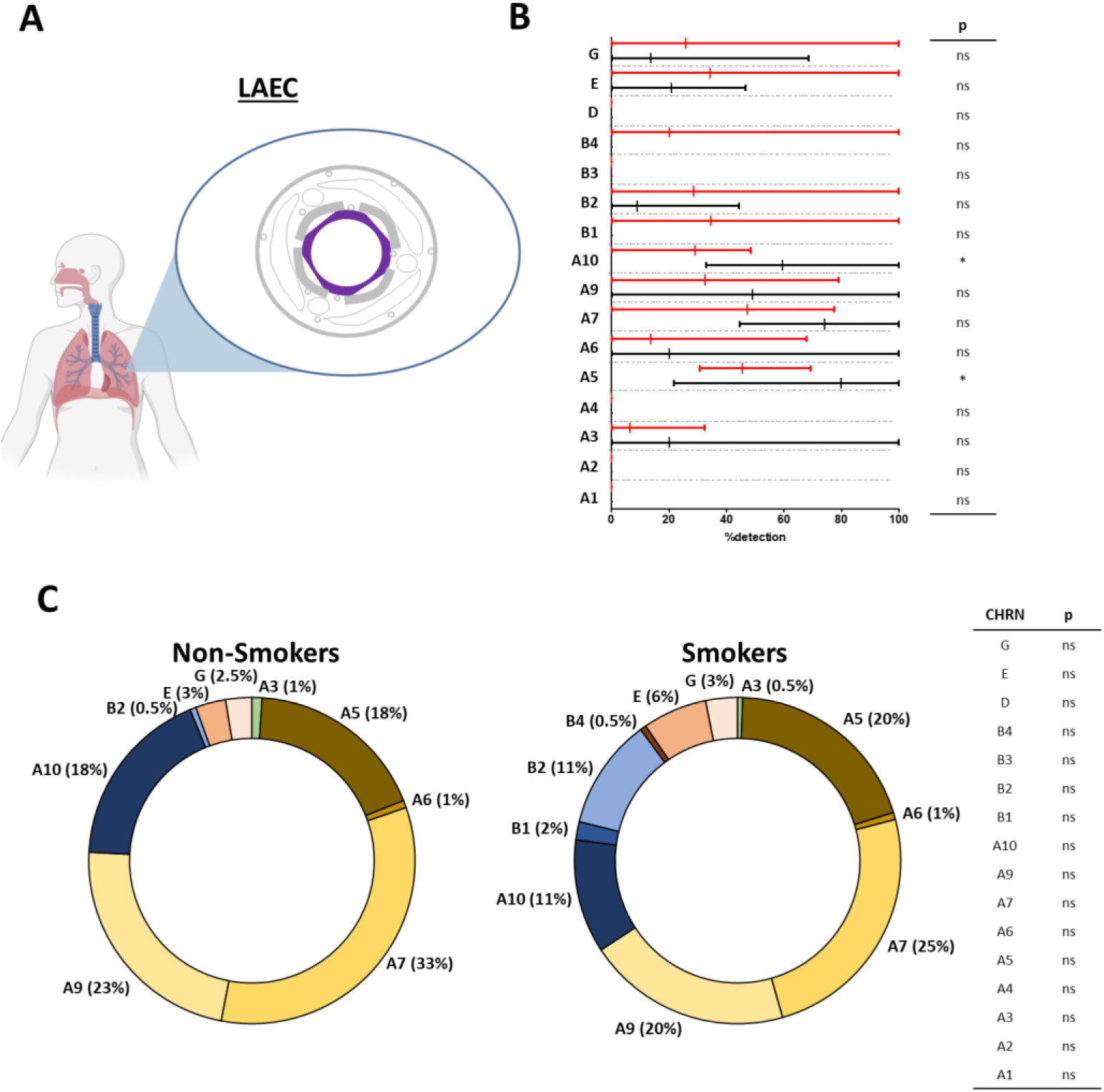
Evaluation of nAChR transcript levels in human large airway epithelial cells (LAEC). **(A)**. Illustrations depicting the origin of the samples. Isolated AEC were collected from bronchi. Large airway epithelial cells (LAEC) are depicted in purple. **(B)**. Histograms showing the detection of nAChRs in non-smokers (black) and smokers (red). **(C)**. Pie charts showing the repartition of nAChRs in non-smokers (left) and smokers (right). *p<0.05 non-smokers (n=5) vs smokers (n=5).

In small AEC (SAEC) (***Figure 3A***), the 16 nAChRs were detected among non-smokers with moderate or low expressions (***Figure 3B***). There was a significant increase of CHRNA5/A7/B2/B3 transcript levels in smokers compared to non-smokers. The global repartition in non-smokers and smokers favoured CHRNA7/A9/A10/B2 representing half of nAChR expressed in SAEC (***Figure 3C***). There was a significant increase of CHRNA7/B2/B3 and a significant decrease of CHRNA2/A9 in the repartition of nAChRs in smokers compared to non-smokers.

**Figure 3.**
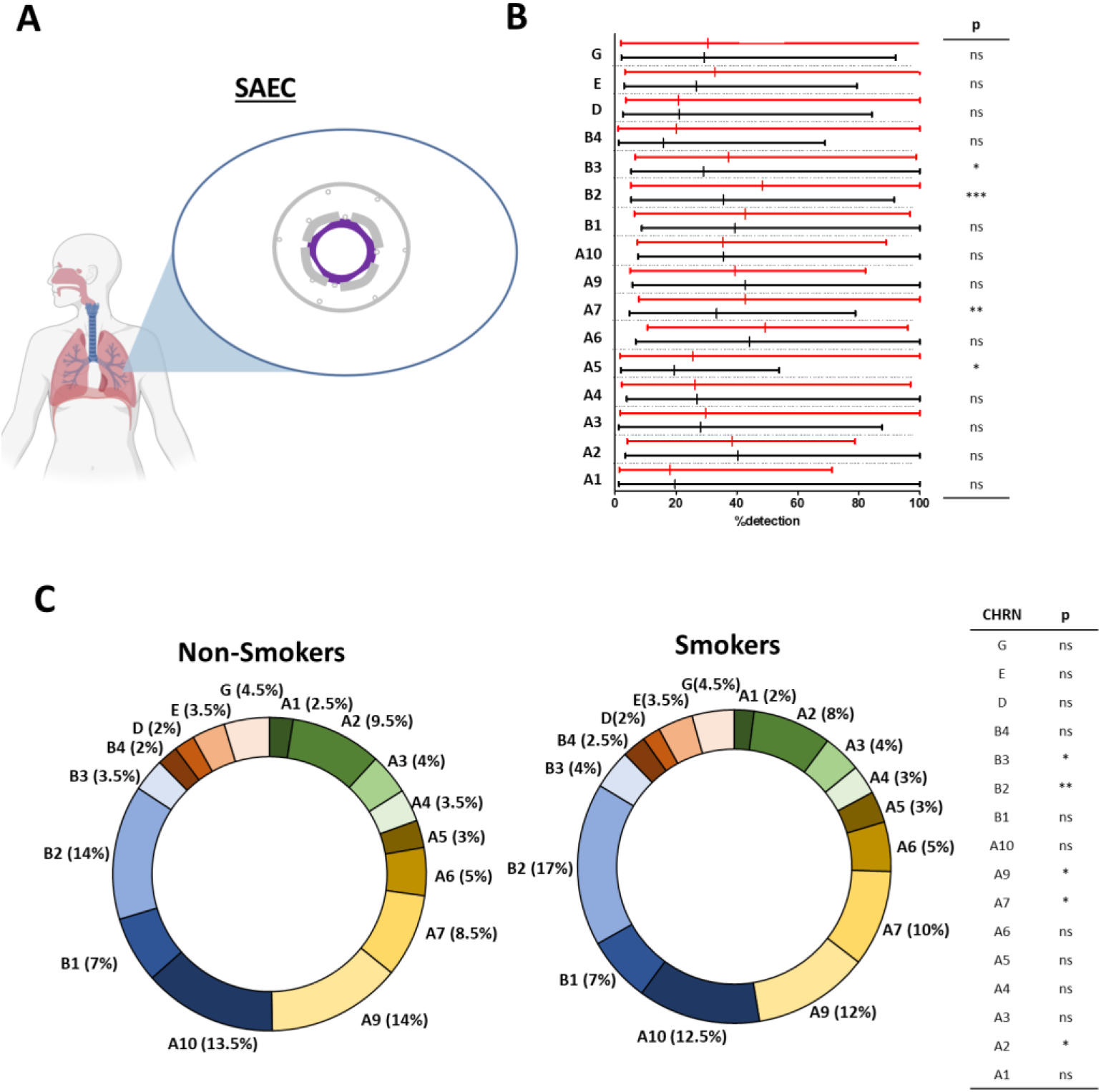
Evaluation of nAChR transcript levels in human small airway epithelial cells (SAEC). **(A)**. Illustrations depicting the origin of the samples. Isolated AEC were collected from bronchioles. Small airway epithelial cells (SAEC) are depicted in purple. **(B)**. Histograms showing the detection of nAChRs in non-smokers (black) and smokers (red). **(C)**. Pie charts showing the repartition of nAChRs in non-smokers (left) and smokers (right). *p<0.05; **p<0.01; ***p<0.001 non-smokers (n=63) vs smokers (n=72).

### Differential pulmonary nAChRs transcripts expressions at a single cell scale

At the level of single cell transcriptomes (***Figure 4***), CHRNA5/A7/A9/A10/B1/E were highly expressed in most of the AEC populations including alveolar, basal, goblet, multiciliated and club cells. CHRNA1/A2/A4/A6/B2/B3/D/G were showing low to no expression in AEC. Interestingly, functional AEC cell populations were distinguished with their nAChR signatures: pneumocytes expressed CHRNA5/A10/B1; basal cells expressed CHRNA5/A7/A10/B1/E; goblet cells expressed CHRNA7/A10/B1/E; multiciliated cells expressed CHRNA9/10/B1/E; club cells expressed CHRNA7/A10/B1; and ionocytes expressed CHRNA3/B4/E.

**Figure 4.**
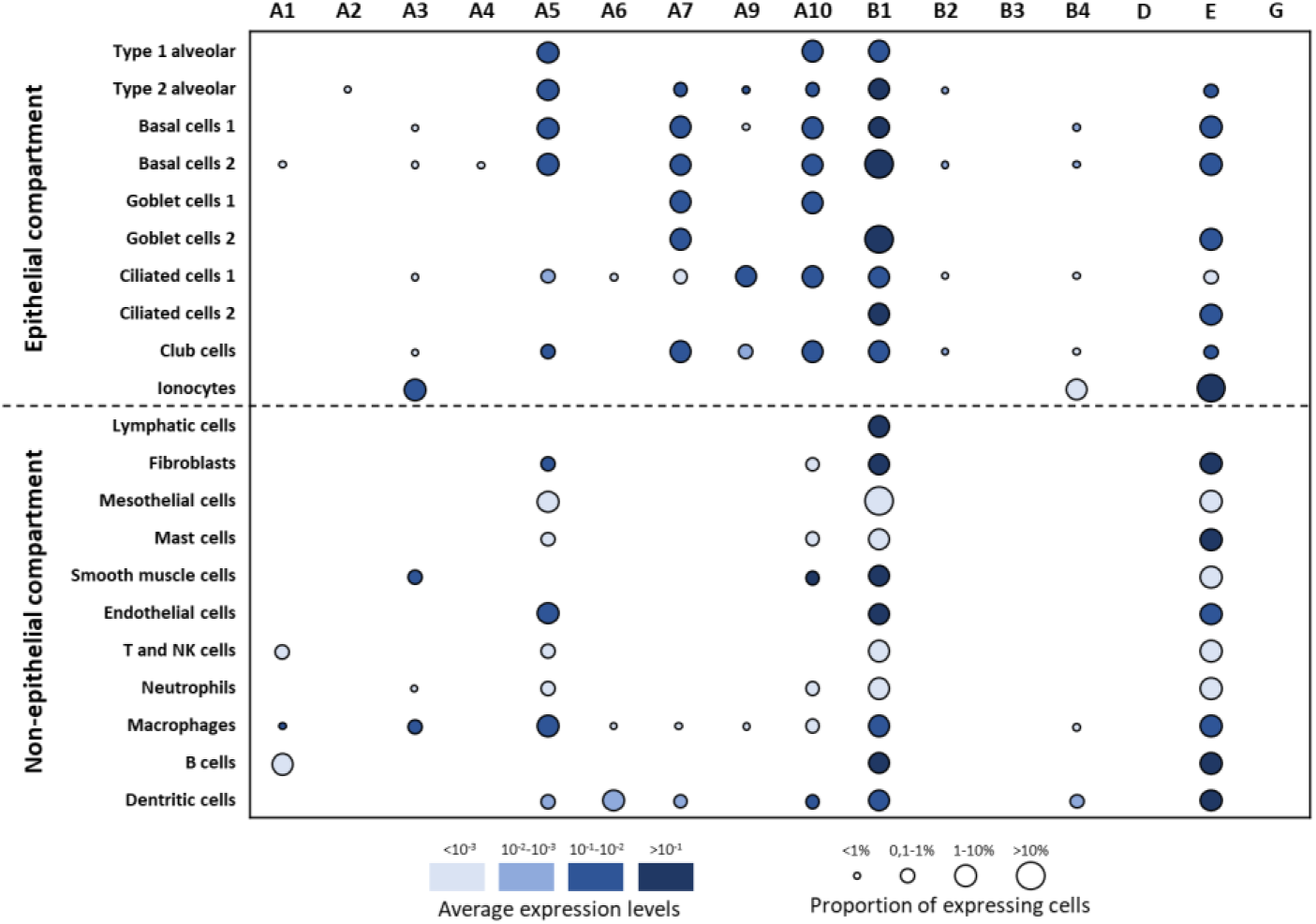
Evaluation of nAChR expressions in lung single cell populations. Dot plots of nAChR expressions in the epithelial and non-epithelial compartments. The identities of cell populations are shown on the y-axis, and the subunits on the x-axis. The colour intensity represents the average expression level, and the size of the dots represents the proportion of the expressing cells in each population. Raw expression values were normalized, log transformed and summarized by published cell clustering.

Regarding non-epithelial cells, CHRNA1/A3/A5/A10/B1/E were highly expressed in specific populations of immune cells including macrophages, B cells, dendritic cells and mast cells. CHRNB1 expression was specific of lymphatic cells. CHRNA5/B1/E were highly expressed in fibroblasts; CHRNA3/10/B1 in smooth muscle cells; CHRNA5/B1/E in endothelial cells and macrophages. B cells and dendritic cells mainly expressed CHRNB1/E.

### Identification of nAChRs in bronchial and bronchiolar epithelia

To investigate nAChR localization in the lung, we selected commercially available validated primary antibodies displaying the antigenic sequences demonstrating the lowest percentage of identity with regard to cross-reactivity (***Supplemental Table S2 and S3, Supplemental Figure S1***). We focussed here on bronchi and large bronchioles as well as smooth muscle and blood vessels on FFPE tissues (***Figure 5A and B***). CHRNA1/A2/A4/B3/G were not detected. CHRNA3 seemed restricted to the apical side of differentiated cells. Surprisingly, CHRNA5 was systematically found in AEC nuclei and the apical side of differentiated AEC, while its pattern was consistent with membrane-bound receptors on smooth muscle cells. CHRNA6/A9 presented similar staining in differentiated AEC, such as CHRNA7/A10/B1/B2/D/E which in addition were found in non-differentiated AEC. Finally, CHRNB4 appeared in multiciliated cells only. When available, our observations were generally concordant with the data from the Human Protein Atlas (***Figure 5B and Supplemental Figure S2***).

**Figure 5.**
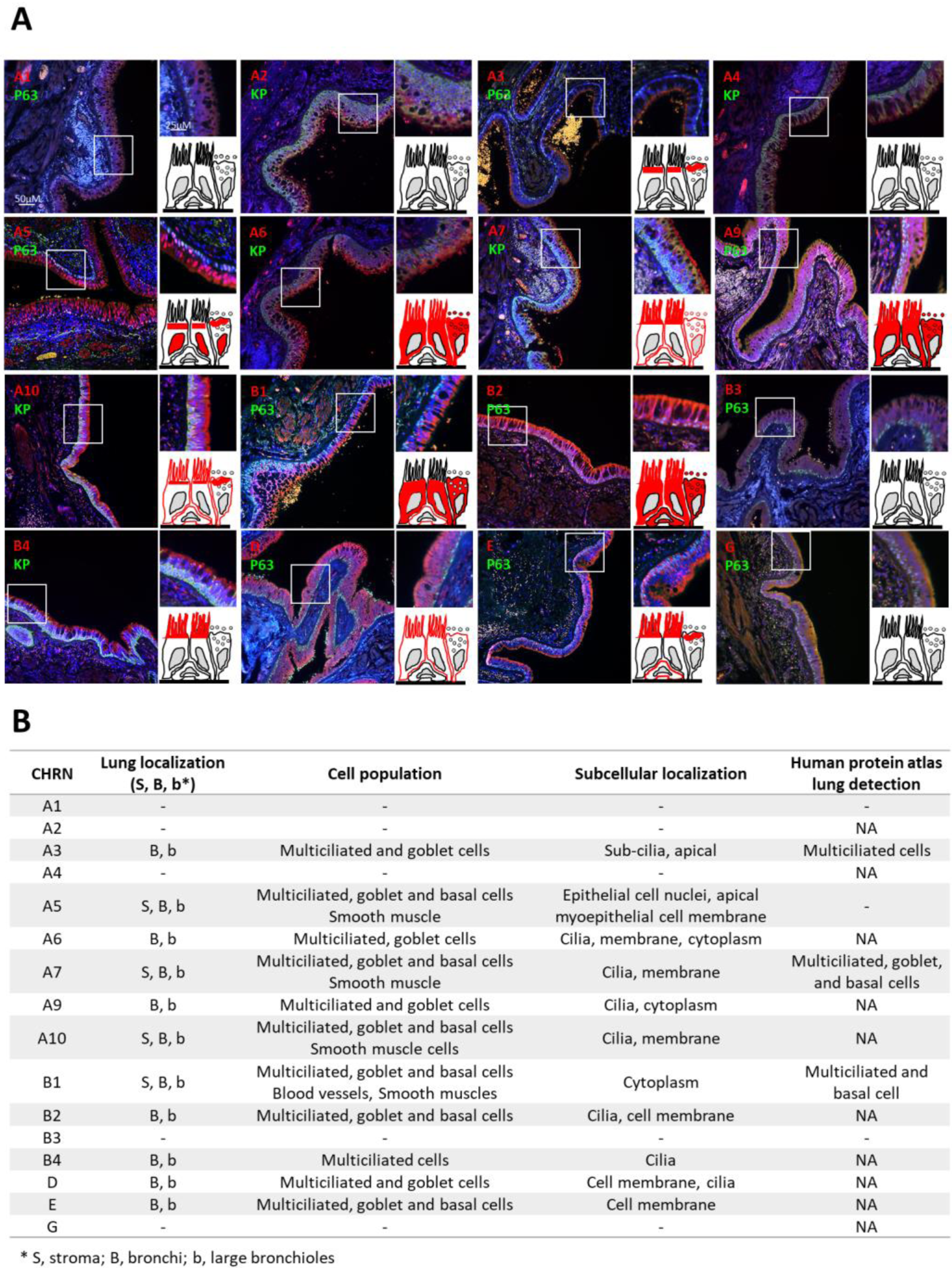
nAChR localizations in human respiratory epithelia. **(A)**. Representative micrographs showing the bronchial epithelia on FFPE lung tissues stained for the nAChRs (all red); non-differentiated cells (p63 or pan-cytokeratin, green) and cell nuclei (DAPI, blue). Magnification corresponding to the selected area is shown. Cartoons depict the localization of each nAChR (in red). **(B)**. Table summarizing nAChR cellular and sub-cellular localization, and the available microscopic data from the Human Protein Atlas (https://www.proteinatlas.org/). NA, not available; -, no detection.

## DISCUSSION

This is the first study showing transcript levels and localizations of all nAChRs in the human adult lung. Interestingly, we identified distinct variations in terms of nAChR transcript levels between whole lung tissues, LAEC and SAEC, as well as important changes between non-smokers and smokers. Since whole lung transcriptomics encompasses all pulmonary tissues, isolated cell studies represent the ideal strategy to unveil nAChR functions in airways. It has been successfully implemented in the context of AEC differentiation analysis, asthma and idiopathic pulmonary fibrosis [27–31]. As such, our comparative analysis points mainly towards CHRNA5/A7/A10/B2/B3/E to tackle the association of nAChR expression and smoking. Furthermore, the impact of smoking could be tie into the associated risks of respiratory diseases, provided complete clinical data are available.

We included in the analysis of nAChR expression levels 298 subjects in three distinct modalities (whole lung tissues, LAEC and SAEC) and performed a preliminary identification of single cell transcriptomic signature. Our immunostaining analyses provide important data regarding the sub-cellular localizations of nAChRs in bronchi and large bronchioles. Microscopic observations and transcriptomic analysis were generally concordant. Because of their modalities of association at the cell membrane and their high degree of amino acids identity, nAChRs immunostainings are generally sparse, rarely concordant, and performed on murine tissues in the literature [32–35]. Nonetheless, since we selected all our antibodies based on thorough sequence alignments of the antigenic sequences, we provide here a complete and robust description of all nAChRs in bronchi and large bronchioles.

Additional studies on larger cohorts are needed to complement and refine our analysis. Deciphering the cellular and molecular impact of the observed differences in transcript expressions in the context of smoking will be essential to understand nAChR-associated pathogenesis. It will be particularly insightful for at least three lung diseases where smoking may partly impact homeostasis: lung cancers, COPD, and COVID-19. (i) nAChRs single nucleotide polymorphisms (SNP) were associated to lung cancer cells [36,37] and nAChRs were shown to be involved in cancer cell proliferation and survival [38–41]. (ii) nAChRs SNPs were also associated with nicotine dependence and COPD [42–46]. (iii) Nicotine, the main ligand of nAChRs, and more generally smoking, have been shown *in vitro* and *in vivo* to modulate the expression of hACE2, the main receptor of SARS-CoV-2 spike S protein [47–50]. In addition, nicotine impairs the renin-angiotensin system (RAS) homeostasis [21], so nicotine-induced imbalance of RAS via nAChRs is a strong candidate to understand hypertension, dysregulated cardiac remodelling, vascular dysfunction and chronic lung diseases associated with severe forms of COVID-19. Interestingly, CHRNA7 transcripts were correlated with hACE2 levels [51] feeding the possible connections between hACE2 and nAChRs localizations in airways [52–54].

We provide the first atlas of nAChRs in the lung and invite cartographers to complete the map in order to provide fundamental understanding of these crucial actors of homeostasis that may contribute to chronic and acute respiratory diseases.

## List of nonstandard abbreviations

(nAChRs): Nicotinic acetylcholine receptors
(AEC): Airway epithelial cells
(COPD): Chronic obstructive pulmonary disease
(COVID-19): Coronavirus disease 2019
(ACE-2): Angiotensin converting enzyme-2
(SARS-CoV-2): Severe acute respiratory syndrome coronavirus
(FFPE): Formalin-fixed paraffin-embedded
(LAEC): Large airway epithelial cell
(SAEC): Small airway epithelial cell
(SNP): Single nucleotide polymorphisms
(RAS): Renin-angiotensin system

## Acknowledgments

We thank the members of the Inserm UMR-S 1250 unit and our collaborators for their helpful comments and insights. We thank the Platform of Cell and Tissue Imaging (PICT) for technical assistance.

## Contributors

ZD performed experiments, analysed the data and wrote the manuscript; JMP collected samples and analysed the data; RB performed experiment; JA, PM and GoD collected samples; MP, PG and UM contributed to experimental analysis and interpretation of the results; GaD collected samples, analysed the data and supervised the experiments; VD designed the experiments, analysed the data and wrote the manuscript. All the authors contributed to the writing and critical appraisal of the manuscript.

## Funding

This work was supported by Funding from University of Reims Champagne-Ardenne (URCA), the French National Institute of Health and Medical Research (Inserm) and a grant from the Research Institute in Public Health (IReSP) in association with the National Institute of Cancer (INCa). It was carried out in the framework of the Federative Research Structure *CAP-Santé* and benefited from the Project Research and Innovation in Inflammatory Respiratory Diseases (RINNOPARI).

## Competing Interests

Dr. Deslée reports personal fees from Nuvaira, personal fees from BTG/PneumRx, personal fees from Chiesi, personal fees from Boehringer, personal fees from Astra Zeneca, outside the submitted work. Dr. Dormoy reports personal fees from Chiesi outside the submitted work.

## Ethics approval

Subjects were recruited from the Department of pulmonary medicine at university hospital of Reims (France) and included in the cohort for Research and Innovation in Chronic Inflammatory Respiratory Diseases (RINNOPARI, NCT02924818). The study was approved by the ethics committee for the protection of human beings involved in biomedical research (CCP Dijon EST I, N°2016-A00242-49) and was conducted in accordance with the ethical guideline of Declaration of Helsinki.

## Notes

### Competing Interest Statement

Dr. Deslee reports personal fees from Nuvaira, personal fees from BTG/PneumRx, personal fees from Chiesi, personal fees from Boehringer, personal fees from Astra Zeneca, outside the submitted work. Dr. Dormoy reports personal fees from Chiesi outside the submitted work.

