## Supplemental for "Nicotinic receptors atlas in the adult human lung"

1. **Supporting information Tables**

**Table S1. List of primers.**

| GENES | GenBank | Forward sequence | Reverse sequence |
|---|---|---|---|
| **CHRNA1** | NM_001039523.2 | 5'- GTCCACACAAGCTCCGGTA-3' | 5'- CAGACGGGTCTCATGTTCG-3' |
| **CHRNA2** | NM_000742.3 | 5'- CTGTGGTGGCTCCTTCTGA-3' | 5'- GGGAGAGGAGAGTGGGTCTC-3' |
| **CHRNA3** | NM_000743.4 | 5'- TGAAATGGAACCCCTCTGAC-3' | 5'- GAAATCCCCAACAGCATTGT-3' |
| **CHRNA4** | NM_000744.6 | 5'- GCCGGACATCGTCCTCTAC-3' | 5'- TGCAGGAGCTCTTGTAAATGG-3' |
| **CHRNA5** | NM_000745.3 | 5'- GACAACAAACGTCTGGTTGAAA-3' | 5'- ACAGAGTCTGAAGGAACACGTATAAC-3' |
| **CHRNA6** | NM_004198.3 | 5'- TTCATGGGGGCTTGTGTC-3' | 5'- GAGCCTCTCCTCAGTTGCAC-3' |
| **CHRNA7** | NM_000746.5 | 5'- CAATGACTCGCAACCACTCA-3' | 5'- GTGATCTGTCCAAGACATTTGC-3' |
| **CHRNA9** | NM_017581.3 | 5'-TCAGAAAATGTGCCCCTGAT-3' | 5'- GGCCCCACAGAAGTGGATA-3' |
| **CHRNA10** | NM_020402.3 | 5'- CCCAGATCATCGACATGGA-3' | 5'- CCCATCGTAGGTAGGCATCT-3' |
| **CHRNB1** | NM_000747.2 | 5'- CACAAAGGTGTACTTAGACCTGGA-3' | 5'- TTCAGTAGCACCACGTCAGG-3' |
| **CHRNB2** | NM_000748.2 | 5'- CTGGCCCAGCTCATCAGT-3' | 5'- TCCAGGTGAGGCGATAATCT-3' |
| **CHRNB3** | NM_000749.4 | 5'- GGTCCGCCCTGTATTACATTC-3' | 5'- TCAGCTGATTCTTTTCATCCAC-3' |
| **CHRNB4** | NM_000750.4 | 5'- TGACGATGAAGACCAGAGTGTC-3' | 5'- GGACGCACACAAACATGAAC-3' |
| **CHRND** | NM_000751.3 | 5'- GGGACCAGAACAATTACAATGAG-3' | 5'- GCAGGAAGATCCAGGCTGT-3' |
| **CHRNE** | NM_000080.4 | 5'- CGACACAGAGGCCTATACTGAG-3' | 5'- GCGGATGATGAGCGAGTAG-3' |
| **CHRNG** | NM_005199.4 | 5'- AGCAGAGTCACTTTGACAATGG-3' | 5'- GTAGTGGGCCATGAGGAAGA-3' |

**Table S2. List of CHRN antibodies.**

| Antibodies | Species | Reference | Companies | Concentrations |
|---|---|---|---|---|
| **α1** | Rabbit | HPA071554 | Sigma-Aldrich | 1:100 |
| **α2** | Mouse | NBP2-61667 | Novus Biological | 1:50 |
| **α3** | Rabbit | HPA029430 | Sigma-Aldrich | 1:100 |
| **α4** | Mouse | NBP2-61674 | Novus Biological | 1:100 |
| **α5** | Rabbit | HPA054381 | Sigma-Aldrich | 1:50 |
| **α6** | Mouse | NBP2-61679 | Novus Biological | 1:100 |
| **α7** | Mouse | NBP2-61738 | Novus Biological | 1:100 |
| **α9** | Rabbit | 26025-1-AP | Proteintech | 1:100 |
| **α10** | Mouse | NBP2-61666 | Novus Biological | 1:50 |
| **β1** | Rabbit | HPA005822 | Sigma-Aldrich | 1:100 |
| **β2** | Rabbit | 17844-1-AP | Proteintech | 1:50 |
| **β3** | Rabbit | APrEST84413 | Novus Biological | 1:100 |
| **β4** | Mouse | NBP2-61742 | Novus Biological | 1:100 |
| **δ** | Rabbit | HPA056404 | Sigma-Aldrich | 1:100 |
| **ε** | Rabbit | NBP1-79951 | Novus Biological | 1:100 |
| **γ** | Rabbit | NBP1-79952 | Novus Biological | 1:100 |

**Table S3. List of recognition antigens of CHRN antibodies and their percentages of identity.**

| s.u. | Antigenic sequences | Position | Identity* |
|---|---|---|---|
| **α1** | MKLGTWTYDGSVVAINPESDQPDLSNFMESGEWVIKESRGWKHSVTYSCCPDTPYLDITYHF | 189-250 | α2/3/4/6 |
| **α2** | EEAKRPPPRAPGDPLSSPSPTALPQGGSHTETEDRLFKHLFRGYNRWARPVPNTSDVVIVRFGLSIAQLIDVDEKNQMMTTNVWLKQEWSDYKLRWNPTDFGNITSLRVPSEMIWIPDIVLYNNADGEFAVTHMTKAHLFSTGTVHWVPPAIYKSSCSIDVTFFPFDQQNCKMKFGSWTYDKAKIDLEQMEQTVDLKDYWESGEWAIVNATGTYNSKKYDCCAEIYPDVTYAFVIRRL | 27-264 | α4<br>α3/5/6<br>β3 |
| **α3** | RTPTTHTMPSWVKTVFLNLLPRVMFMTRPTSNEGNAQKPRPLYGAELSNLNCFSRAESKGCKEGYPCQDGMCGYCHHRRIKISNFSANLTRSSSSESVDAVLSLSALSPEIKEAIQSVKYIAENMKAQNEAKEIQDDWKYVAM | 331-473 | α1/2/6<br>β2/4 |
| **α4** | HVETRAHAEERLLKKLFSGYNKWSRPVANISDVVLVRFGLSIAQLIDVDEKNQMMTTNVWVKQEWHDYKLRWDPADYENVTSIRIPSELIWRPDIVLYNNADGDFAVTHLTKAHLFHDGRVQWTPPAIYKSSCSIDVTFFPFDQQNCTMKFGSWTYDKAKIDLVNMHSRVDQLDFWESGEWVIVDAVGTYNTRKYECCAEIYPDITYAFVIRRL | 29-242 | α2<br>α3/5/6<br>β3 |
| **α5** | AQRGLSEPSSIAKHEDSLLKDLFQDYERWVRPVEHLNDK | 33-71 | β3<br>α4 |
| **α6** | KGCVGCATEERLFHKLFSHYNQFIRPVENVSDPVTVHFEVAITQLANVDEVNQIMETNLWLRHIWNDYKLRWDPMEYDGIETLRVPADKIWKPDIVLYNNAVGDFQVEGKTKALLKYNGMITWTPPAIFKSSCPMDITFFPFDHQNCSLKFGSWTYDKAEIDLLIIGSKVDMNDFWENSEWEIIDASGYKHDIKYNCCEEIYTDITYSFYIRRL | 26-239 | α3<br>α2/4/5<br>β3 |
| **α7** | LYKELVKNYNPLERPVANDSQPLTVYFSLSLLQIMDVDEKNQVLTTNIWLQMSWTDHYLQWNVSEYPGVKTVRFPDGQIWKPDILLYNSADE | 52-259 | α2/3/4/10 |
| **α9** | HFCGAEARPVPHWARVVILKYMSRVLFVYDVGESCLSPHHSRERDHLTKVYSKLPESNLKAARNKDLSRKKDMNKRLKNDLGCQGKNPQEAESYCAQYKVLTRNIEYIAKCLKDHKATNSKGSEWKKVAKVIDRFFMWIFFIMVFVMTILIIARAD | 324-479 | α2/4/6/10 |
| **α10** | AEGRLALKLFRDLFANYTSALRPVADTDQTLNVTLEVTLSQIIDMDERNQVLTLYLWIRQEWTDAYLRWDPNAYGGLDAIRIPSSLVWRPDIVLYNKADAQPPGSASTNVVLRHDGAVRWDAPAITRSSCRVDVAAFPFDAQHCGLTFGSWTHGGHQLDVRPRGAAASLADFVENVEWRVLGMPARRRVLTYGCCSEPYPDVTFTLLLRRRAA | 25-237 | α9<br>α3/7 |
| **β1** | SVCVLNLHHRSPHTHQMPLWVRQIFIHKLPLYLRLKRPKPERDLMPEPPHCSSPGSGWGRGTDEYFIRKPPSDFLfpkpNRFQPELSAPDLRRFIDGPNRAVALLPELREVVSSISYIARQLQEQEDHDALKEDWQF | 314-463 | α3<br>β2/4 |
| **β2** | LLRLCSGVWGTDTEERLVEHLLDPsrYNKLIRPATNGSELVTVQLMVSLAQLISVHEREQIMTTNVWLTQEWEDYRLTWKPEEFDNMKKVRLPSKHIWLPDVVLYNNADGMYEVSFYSNAVVSYDGSIFWLPPAIYKSACKIEVKHFPFDQQNCTMKFRSWTYDRTEIDLVLKSEVASLDDFTPSGEWDIVALPGRRNENPDDSTYVDITYD | 16-227 | β4<br>α2 |
| **β3** | TGFNSIAENEDALLRHLFQGYQKWVRPVLHSNDTI | 20-54 | α5<br>α2/4 |
| **β4** | RVANAEEKLMDDLLNKtrYNNLIRPATSSSQLISIKLQLSLAQLISVNEREQIMTTNVWLKQEWTDYRLTWNSSRYEGVNILRIPAKRIWLPDIVLYNNADGTYEVSVYTNLIVRSNGSVLWLPPAIYKSACKIEVKYFPFDQQNCTLKFRSWTYDHTEIDMVLMTPTASMDDFTPSGEWDIVALPGRRTVNPQDPSYVDVTYDFIIKRKPLFYT | 22-236 | β2<br>α3<br>β3 |
| **δ** | LVRRSSSLGYISKAEEYFLLKSRSDLMfekqserhglARRLTTARRPPASSEQAQQELFNELKPAVDGANFIVNHMRDQNNYNEEKDSWNR | 373-464 | β2 |
| **ε** | GLLGRGVGKNEELRLYHHLFNNYDPGSRPVREPEDTVTISLKVTLTNLIS | 13-62 | α6, γ |
| **γ** | NYDPNLRPAERDSDVVNVSLKLTLTNLISLNEREEALTTNVWIEMQWCDY | 36-85 | δ<br>α7, β2/4 |

*Range of the percentage of identity obtained from blastp: yellow, 65-80%; light green, 50-65%; dark green, <50%

### 3. Supporting information Figures

**Figure S1. Constraint-based Multiple Alignment of CHRN antibodies.** COBALT alignment is shown for the 16 subunits and antigen sequences of corresponding antibodies are highlighted in grey. Red amino acids are conserved for all subunits.

```
α1    001  MEPW--------PLLLL-------FSLCS--------AGL-------VLGSEH------------ETRLVAKLFKD--YS  36
α2    001  MGPS-cpvflSFTKLSLwwllLTPAGGEEakr-ppprAPGdplsspsPTALPQggshte-t----EDRLFKHLFRG--YN  71
α3    001  M----gsgplSLPLALSpprlLLLLLLSL--------LPV-------ARASEA------------EHRLFERLFED--YN  47
α4    001  M----------------------ELGGPGapr-llp-----plllllGTGLLRasshve-trahaEERLLKKLFSG--YN  49
α5    001  MAAR-gsgprALRLLLL---vQLVAGRCG--------LAG-------AAGGAQrglsepssiakhEDSLLKDLFQD--YE  59
α6    001  MLTSkgqgflHGGLCL-----WLCVFTPF--------FKG-------CVGCAT------------EERLFHKLFSH--YN  46
α7    001  MRCSpggvwlALAASLLhg--KATASPPStppwdpghIPG-------ASVRPApgpvsl--qgefQRKLYKELVKN--YN  67
α9    001  ---------------MNwshsCISF-------------CW-------IYFAASrlraaetadgkyAQKLFNDLFED--YS  43
α10   001  ---------------MGlrshHLSLGLLLlfl-lpaeCLG-------AEGRLAl-------------KLFRDLFAN--YT  42
β1    001  MTPG--------ALLMLlg----ALGAPL--------APG-------VRGSEA------------EGRLREKLFSG--YD  39
β2    001  MARRcgpvalLLGFGL--------LRLCS----------G-------VWGTDT------------EERLVEHLLDPsrYN  43
β3    001  M----------LPDFML---vLIVLG---------------------IPSSATtgfn---siaenEDALLRHLFQG--YQ  41
β4    001  MRR--apslvL--FFL--------VALCG--------RGN-------CRVANA------------EEKLMDDLLNKtrYN  41
γ     001  MHGG------QGPLLLL-----LLLAVCL----------G-------AQGRNQ------------EERLLADLMQN--YD  38
ε     001  MARA------PLGVLLLl---------------------G-------LLGRGV-gkne-------ELRLYHHLFNN--YD  36
δ     001  MEGP------VLTLGLL-----AALAVC-----------G-------SWGLNE------------EERLIRHLFQEkgYN  39

α1    037  SVVRPVEDHRQVVEVTVGLQLIQLINVDEVNQIVTTNVRLKQgdmvdlprpscvtlgvplfshlqneQWVDYNLKWNPDD  116
α2    072  RWARPVPNTSDVVIVRFGLSIAQLIDVDEKNQMMTTNVWLKQ-------------------------EWSDYKLRWNPTD  126
α3    048  EIIRPVANVSDPVIIHFEVSMSQLVKVDEVNQIMETNLWLKQ-------------------------IWNDYKLKWNPSD  102
α4    050  KWSRPVANISDVVLVRFGLSIAQLIDVDEKNQMMTTNVWVKQ-------------------------EWHDYKLRWDPAD  104
α5    060  RWVRPVEHLNDKIKIKFGLAISQLVDVDEKNQLMTTNVWLKQ-------------------------EWIDVKLRWNPDD  114
α6    047  QFIRPVENVSDPVTVHFEVAITQLANVDEVNQIMETNLWLRH-------------------------IWNDYKLRWDPME  101
α7    068  PLERPVANDSQPLTVYFSLSLLQIMDVDEKNQVLTTNIWLQM-------------------------SWTDHYLQWNVSE  122
α9    044  NALRPVEDTDKVLNVTLQITLSQIKDMDERNQILTAYLWIRQ-------------------------IWHDAYLTWDRDQ  98
α10   043  SALRPVADTDQTLNVTLEVTLSQIIDMDERNQVLTLYLWIRQ-------------------------EWTDAYLRWDPNA  97
β1    040  SSVRPAREVGDRVRVSVGLILAQLISLNEKDEEMSTKVYLDL-------------------------EWTDYRLSWDPAE  94
β2    044  KLIRPATNGSELVTVQLMVSLAQLISVHEREQIMTTNVWLTQ-------------------------EWEDYRLTWKPEE  98
β3    042  KWVRPVLHSNDTIKVYFGLKISQLVDVDEKNQLMTTNVWLKQ-------------------------EWTDHKLRWNPDD  96
β4    042  NLIRPATSSSQLISIKLQLSLAQLISVNEREQIMTTNVWLKQ-------------------------EWTDYRLTWNSSR  96
γ     039  PNLRPAERDSDVVNVSLKLTLTNLISLNEREEALTTNVWIEM-------------------------QWCDYRLRWDPRD  93
ε     037  PGSRPVREPEDTVTISLKVTLTNLISLNEKEETLTTSVWIGI-------------------------DWQDYRLNYSKDD  91
δ     040  KELRPVAHKEESVDVALALTLSNLISLKEVEETLTTNVWIEH-------------------------GWTDNRLKWNAEE  94

α1    117  YGGVKKIHIPSEKIWRPDLVLYNNADGDFAIVKFTKVLLQYTGHITWTPPAIFKSYCEIIVTHFPFDEQNCSMKLGTWTY  196
α2    127  FGNITSLRVPSEMIWIPDIVLYNNADGEFAVTHMTKAHLFSTGTVHWVPPAIYKSSCSIDVTFFPFDQQNCKMKFGSWTY  206
α3    103  YGGAEFMRVPAQKIWKPDIVLYNNAVGDFQVDDKTKALLKYTGEVTWIPPAIFKSSCKIDVTYFPFDYQNCTMKFGSWSY  182
α4    105  YENVTSIRIPSELIWRPDIVLYNNADGDFAVTHLTKAHLFHDGRVQWTPPAIYKSSCSIDVTFFPFDQQNCTMKFGSWTY  184
α5    115  YGGIKVIRVPSDSVWTPDIVLFDNADGRFEGT-STKTVIRYNGTVTWTPPANYKSSCTIDVTFFPFDLQNCSMKFGSWTY  193
α6    102  YDGIETLRVPADKIWKPDIVLYNNAVGDFQVEGKTKALLKYNGMITWTPPAIFKSSCPMDITFFPFDHQNCSLKFGSWTY  181
α7    123  YPGVKTVRFPDGQIWKPDILLYNSADERFDATFHTNVLVNSSGHCQYLPPGIFKSSCYIDVRWFPFDVQHCKLKFGSWSY  202
α9    99   YDGLDSIRIPSDLVWRPDIVLYNKADDESSEPVNTNVVLRYDGLITWDAPAITKSSCVVDVTYFPFDNQQCNLTFGSWTY  178
α10   98   YGGLDAIRIPSSLVWRPDIVLYNKADAQPPGSASTNVVLRHDGAVRWDAPAITRSSCRVDVAAFPFDAQHCGLTFGSWTH  177
β1    95   HDGIDSLRITAESVWLPDVVLLNNNDGNFDVALDISVVVSSDGSVRWQPPGIYRSSCSIQVTYFPFDWQNCTMVFSSYSY  174
β2    99   FDNMKKVRLPSKHIWLPDVVLYNNADGMYEVSFYSNAVVSYDGSIFWLPPAIYKSACKIEVKHFPFDQQNCTMKFRSWTY  178
β3    97   YGGIHSIKVPSESLWLPDIVLFENADGRFEGSLMTKVIVKSNGTVVWTPPASYKSSCTMDVTFFPFDRQNCSMKFGSWTY  176
β4    97   YEGVNILRIPAKRIWLPDIVLYNNADGTYEVSVYTNLIVRSNGSVLWLPPAIYKSACKIEVKYFPFDQQNCTLKFRSWTY  176
γ     94   YEGLWVLRVPSTMVWRPDIVLENNVDGVFEVALYCNVLVSPDGCIYWLPPAIFRSACSISVTYFPFDWQNCSLIFQSQTY  173
ε     92   FGGIETLRVPSELVWLPEIVLENNIDGQFGVAYDANVLVYEGGSVTWLPPAIYRSVCAVEVTYFPFDWQNCSLIFRSQTY  171
δ     95   FGNISVLRLPPDMVWLPEIVLENNNDGSFQISYSCNVLVYHYGFVYWLPPAIFRSSCPISVTYFPFDWQNCSLKFSSLKY  174

α1    197  DGSVVAINPESDQP------------DLSNFMESGEWVIKESRGWK---HSVTYSCCPDTpYLDITYHFVMQRLPLYFIV  261
α2    207  DKAKIDLEQMEQTV------------DLKDYWESGEWAIVNATGTYNS---KKYDCCAEI-YPDVTYAFVIRRLPLFYTI  270
α3    183  DKAKIDLVLIGSSM------------NLKDYWESGEWAIIKAPGYK---HDIKYNCCEEI-YPDITYSLYIRRLPLFYTI  246
α4    185  DKAKIDLVNMHSRV------------DQLDFWESGEWVIVDAVGTYNT---RKYECCAEI-YPDITYAFVIRRLPLFYTI  248
α5    194  DGSQVDIILEDQDV------------DKRDFFDNGEWEIVSATGSK---GNRTDSCCW---YPYVTYSFVIKRLPLFYTL  255
α6    182  DKAEIDLLIIGSKV------------DMNDFWENSEWEIIDASGYK---HDIKYNCCEEI-YTDITYSFYIRRLPMFYTI  245
α7    203  GGWSLDLQMQEADI--------------SGYIPNGEWDLVGIPGKRSE---RFYECCKEP-YPDVTFTVTMRRRTLYYGL  264
α9    179  NGNQVDIFNALDSG------------DLSDFIEDVEWEVHGMPAVKNV---ISYGCCSEP-YPDVTFTLLLKRRSSFYIV  242
α10   178  GGHQLDVRPRGAAA------------SLADFVENVEWRVLGMPARRRV---LTYGCCSEP-YPDVTFTLLLRRRAAAYVC  241
β1    175  DSSEVSLQTGLGPDgqghq---eihiHEGTFIENGQWEIIHKPSRLIQpPGDPRGGREGQ-RQEVIFYLIIRRKPLFYLV  250
β2    179  DRTEIDLVLKSEVA------------SLDDFTPSGEWDIVALPGRR---NENPDDST----YVDITYDFIIRRKPLFYTI  239
β3    177  DGTMVDLILINENV------------DRKDFFDNGEWEILNAKGMK---GNRRDGVYS---YPFITYSFVLRRLPLFYTL  238
β4    177  DHTEIDMVLMTPTA------------SMDDFTPSGEWDIVALPGRR---TVNPQDPS----YVDVTYDFIIKRKPLFYTI  237
γ     174  STNEIDLQLSQEDGqt----iewifiDPEAFTENGEWAIQHRPAKMLL---DPAAPAQEAgHQKVVFYLLIQRKPLFYVI  246
ε     172  NAEEVEFTFAVDNDgk---tinkidiDTEAYTENGEWAIDFCPGVIRR---HHGGATDGPgETDVIYSLIIRRKPLFYVI  245
δ     175  TAKEITLSLKQDAKenrtypvewiiiDPEGFTENGEWEIVHRPARVNV---DPRAPLDSPsRQDITFYLIIRRKPLFYII  251
```

```
α1     262  NVIIPCLLFSFLTGLVFYLPTDSG-EKMTLSISVLLSLTVFLLVIVELIPSTSSAVPLIGKYMLFTMVFVIASIIITVIV  340
α2     271  NLIIPCLLISCLTVLVFYLPSDCG-EKITLCISVLLSLTVFLLLITEIIPSTSLVIPLIGEYLLFTMIFVTLSIVITVFV  349
α3     247  NLIIPCLLISFLTVLVFYLPSDCG-EKVTLCISVLLSLTVFLLVITETIPSTSLVIPLIGEYLLFTMIFVTLSIVITVFV  325
α4     249  NLIIPCLLISCLTVLVFYLPSECG-EKITLCISVLLSLTVFLLLITEIIPSTSLVIPLIGEYLLFTMIFVTLSIVITVFV  327
α5     256  FLIIPCIGLSFLTVLVFYLPSNEG-EKICLCTSVLVSLTVFLLVIEEIIPSSSKVIPLIGEYLVFTMIFVTLSIMVTVFA  334
α6     246  NLIIPCLFISFLTVLVFYLPSDCG-EKVTLCISVLLSLTVFLLVITETIPSTSLVVPLVGEYLLFTMIFVTLSIVVTVFV  324
α7     265  NLLIPCVLISALALLVFLLPADSG-EKISLGITVLLSLTVFMLLVAEIMPATSDSVPLIAQYFASTMIIVGLSVVVTVIV  343
α9     243  NLLIPCVLISFLAPLSFYLPAASG-EKVSLGVTILLAMTVFQLMVAEIMPA-SENVPLIGKYYIATMALITASTALTIMV  320
α10    242  NLLLPCVLISLLAPLAFHLPADSG-EKVSLGVTVLLALTVFQLLLAESMPP-AESVPLIGKYYMATMTMVTFSTALTILI  319
β1     251  NVIAPCILITLLAIFVFYLPPDAG-EKMGLSIFALLTLTVFLLLLADKVPETSLSVPIIIKYLMFTMVLVTFSVILSVVV  329
β2     240  NLIIPCVLITSLAILVFYLPSDCG-EKMTLCISVLLALTVFLLLISKIVPPTSLDVPLVGKYLMFTMVLVTFSIVTSVCV  318
β3     239  FLIIPCLGLSFLTVLVFYLPSDEG-EKLSLSTSVLVSLTVFLLVIEEIIPSSSKVIPLIGEYLLFIMIFVTLSIIVTVFV  317
β4     238  NLIIPCVLTTLLAILVFYLPSDCG-EKMTLCISVLLALTFFLLLISKIVPPTSLDVPLIGKYLMFTMVLVTFSIVTSVCV  316
γ      247  NIIAPCVLISSVAILIHFLPAKAGgQKCTVAINVLLAQTVFLFLVAKKVPETSQAVPLISKYLTFLLVVTILIVVNAVVV  326
ε      246  NIIVPCVLISGLVLLAYFLPAQAGgQKCTVSINVLLAQTVFLFLIAQKIPETSLSVPLLGRFLIFVMVVATLIVMNCVIV  325
δ      252  NILVPCVLISFMVNLVFYLPADSG-EKTSVAISVLLAQSVFLLLISKRLPATSMAIPLIGKFLLFGMVLVTMVVVICVIV  330

α1     341  INTHHRSPSTHV-MPNWVRKVFIDTIPNIMFfstMKRPSREKQDKKIFTEDID------------ISDISGKPGP-----  402
α2     350  LNVHHRSPSTHT-MPHWVRGALLGCVPRWLL---MNRP------------------------------------------  383
α3     326  LNVHYRTPTTHT-MPSWVKTVFLNLLPRVMF---MTRPTSNEGNAQKPRPLYG-aELSNLNCFSRAESKGCKEGYPCQD-  399
α4     328  LNVHHRSPRTHT-MPTWVRRVFLDIVPRLLL---MKRPSVVKDNCRRLIESMHkmASAPRFWPEPEGEPPATSGTQSLH-  402
α5     335  INIHHRSSSTHNaMAPLVRKIFLHTLPKLLC---MRSHVDRYFT--------------------QKEETESGSGP-----  386
α6     325  LNIHYRTPTTHT-MPRWVKTVFLKLLPQVLL---MRWP------LDKTRGTGS-dAVPRGLARRPAKGKLASHGEPRHL-  392
α7     344  LQYHHHDPDGGK-MPKWTRVILLNWCAWFLR---MKRPGEDKVRPACQHKQRR--CSLASVEMSAVAPPPASNGNLLYIg  417
α9     321  MNIHFCGAEARP-VPHWARVVILKYMSRVLFvydVGESC-------LSPHHSR-eRDHLTKVYSKLPESNLKAARNKDLs  391
α10    320  MNLHYCGPSVRP-VPAWARALLLGHLARGLCvreRGEPCGQSRPPELSPSPQS-------PEGGAGPPAGPCHEPRCLC-  390
β1     330  LNLHHRSPHTHQ-MPLWVRQIFIHKLPLYLR---LKRPKPERDLMPEPPHCSS----PGSGWGRGTDEYFIRKPPSDFL-  400
β2     319  LNVHHRSPTTHT-MAPWVKVVFLEKLPALLF---MQQPRHHCARQRLRLRRRQ-rEREGAGALFFREAPGADSCT-----  388
β3     318  INVHHRSSSTYHpMAPWVKRLFLQKLPKLLC---MKDHVDRYSSPEKEESQPV----VKGKVLEKKKQKQLSDGE-----  385
β4     317  LNVHHRSPSTHT-MAPWVKRCFLHKLPTFLF---MKRPGPDSSPARAFPPSKScvTKPEATATSTSPSNFYGNSM-----  387
γ      327  LNVSLRSPHTHS-MARGVRKVFLRLLPQLLR---MHVRPLAPAAVQDTQSRLQ--NGSSGWSITTGEEVALCLPRSELL-  399
ε      326  LNVSQRTPTTHA-MSPRLRHVLLELLPRLLG---SPPPPEAPRAASPPRRASS------VGLLLRAEELILKKPRSELV-  394
δ      331  LNIHFRTPSTHV-LSEGVKKLFLETLPELLH---MSRPAEDGPSPGALVRRSS--SLGY---ISKAEEYFLLKSRSDLM-  400

α1     403  ------------------------------PPMGFHSPLIKHPEVKSAIE------------------------------  422
α2     384  ----------------------------------------------------pp--pvel-----------chp------  392
α3     400  ---gmcgychhrrikiSNFSANLTRSSSSESVDAVLSLSALSPEIKEAIQ------------------------------  446
α4     403  -------------------------PPSPSFCVPLDVPAEPGPSCKSPSDqlppqqpleaekasphpspgpcrpphgtqa  457
α5     387  --------------------------------------KSSRNTLEAALD------------------------------  398
α6     393  ---kecfhchk-----SNELATSKRRLSHQPLQWVVENSEHSPEVEDVIN------------------------------  434
α7     418  frgldgvhcvptpdsgVVCGRMACSPTHDEHLLHGGQPPEGDPDLAKILE------------------------------  467
α9     392  rk------------------------------------KDMNKRLKNDLG-cqgknpqeaes------------------  416
α10    391  -------------------------------------------RQEALLH------------------------------  397
β1     401  ---fp---------kpNRFQPELSAPDLRRFIDGPNRAVALLPELREVVS------------------------------  438
β2     389  --------cfvnrasvQGLAGAFGAEPA-PVAGPGRSGEPCGCGLREAVD------------------------------  429
β3     386  --------------------------------------KVLVAFLEKAAD------------------------------  397
β4     388  --------yfvnpasaASKSPAGSTPVAIPRDFWLRSSGRFRQDVQEALE------------------------------  429
γ      400  ---fqq---wqrqglvAAALEKLEKGPELGLSQFCGSLKQAAPAIQACVE------------------------------  443
ε      395  ---feg----------------QRHRQGTWTAAFCQSLGAAAPEVRCCVD------------------------------  425
δ      401  ---fek---qserhglARRLTTARRPP----ASSEQAQQELFNELKPAVD------------------------------  440

α1          --------------------------------------------------------------------------------
α2     393  --lrlklspsyhwlesnvdaeerevvveeedrwacagh--vap-----svgtlcshghlhsg------------------  445
α3          --------------------------------------------------------------------------------
α4     458  pglakarslsvqhmsspgeaveggvrcrsrsiqycvprddaapeadgqaagalasr-nthsaelpppdqpspckctckke  536
α5          --------------------------------------------------------------------------------
α6          --------------------------------------------------------------------------------
α7          --------------------------------------------------------------------------------
α9     417  ---------------------------------ycaqyk-------------vltr------------------------  426
α10         --------------------------------------------------------------------------------
β1          --------------------------------------------------------------------------------
β2          --------------------------------------------------------------------------------
β3          --------------------------------------------------------------------------------
β4          --------------------------------------------------------------------------------
γ           --------------------------------------------------------------------------------
ε           --------------------------------------------------------------------------------
δ           --------------------------------------------------------------------------------
```

```
α1    423  ---------------------------------GIKYIAETMKSDQESNNAAAEWKYVAMVMDHILLGVFMLVCIIGTLA  469
α2    446  --asgpkaeallqege-----lllsphmqkaleGVHYIADHLRSEDADSSVKEDWKYVAMVIDRIFLWLFIIVCFLGTIG  518
α3    447  ---------------------------------SVKYIAENMKAQNEAKEIQDDWKYVAMVIDRIFLWVFTLVCILGTAG  493
α4    537  pssvspsatvktrstkappphlplspaltraveGVQYIADHLKAEDTDFSVKEDWKYVAMVIDRIFLWMFIIVCLLGTVG  616
α5    399  ---------------------------------SIRYITRHIMKENDVREVVEDWKFIAQVLDRMFLWTFLFVSIVGSLG  445
α6    435  ---------------------------------SVQFIAENMKSHNETKEVEDDWKYVAMVVDRVFLWVFIIVCVFGTAG  481
α7    468  ---------------------------------EVRYIANRFRCQDESEAVCSEWKFAACVVDRLCLMAFSVFTIICTIG  514
α9    427  ---------------------------------NIEYIAKCLKDHKATNSKGSEWKKVAKVIDRFFMWIFFIMVFVMTIL  473
α10   398  ---------------------------------HVATIANTFRSHRAAQRCHEDWKRLARVMDRFFLAIFFSMALVMSLL  444
β1    439  ---------------------------------SISYIARQLQEQEDHDALKEDWQFVAMVVDRLFLWTFIIFTSVGTLV  485
β2    430  ---------------------------------GVRFIADHMRSEDDDQSVSEDWKYVAMVIDRLFLWIFVFVCVFGTIG  476
β3    398  ---------------------------------SIRYISRHVKKEHFISQVVQDWKFVAQVLDRIFLWLFLIVSVTGSVL  444
β4    430  ---------------------------------GVSFIAQHMKNDDEDQSVVEDWKYVAMVVDRLFLWVFMFVCVLGTVG  476
γ     444  ---------------------------------ACNLIACARHQQSHFDNGNEEWFLVGRVLDRVCFLAMLSLFICGTAG  490
ε     426  ---------------------------------AVNFVAESTRDQEATGEEVSDWVRMGNALDNICFWAALVLFSVGSSL  472
δ     441  ---------------------------------GANFIVNHMRDQNNYNEEKDSWNRVARTVDRLCLFVVTPVMVVGTAW  487

α1    470  VFAGRLIELNQQG-------------------------------------------------------------------  482
α2    519  LFLPPFLAGMI---------------------------------------------------------------------  529
α3    494  LFLQPLMAREDA--------------------------------------------------------------------  505
α4    617  LFLPPWLAGMI---------------------------------------------------------------------  627
α5    446  LFVPVIYKWANIL---Ipvhignank------------------------------------------------------  468
α6    482  LFLQPLLGNTGKS-------------------------------------------------------------------  494
α7    515  ILMSAPNFVEAVSkdfA---------------------------------------------------------------  531
α9    474  IIARAD--------------------------------------------------------------------------  479
α10   445  VLVQAL--------------------------------------------------------------------------  450
β1    486  IFLDATYHLPPPDpf----------p------------------------------------------------------  501
β2    477  MFLQPLFQNYTTTtflHsdhsapssk------------------------------------------------------  502
β3    445  IFTPALKMW---L---Hsyh------------------------------------------------------------  458
β4    477  LFLPPLFQTHAAS---Egpyaaqrd-------------------------------------------------------  498
γ     491  IFLMAHYNRVPALpfpGdprpy----------------------------------------------------------  512
ε     473  IFLGAYFNRVPDLpyaPciqp-----------------------------------------------------------  493
δ     488  IFLQGVYNQPPPQpfpGdpysynvqdkrfihttpswwwncbinlmnihgovtoolscobaltcobaltcgicmdgetcob  567

α1         ------------
α2         ------------
α3         ------------
α4         ------------
α5         ------------
α6         ------------
α7         ------------
α9         ------------
α10        ------------
β1         ------------
β2         ------------
β3         ------------
β4         ------------
γ     513  -lpspd------  517
ε          ------------
δ     568  altridvbtjjk  579
```

**Figure S2. Localization of nAChRs on lung tissues from the Human Protein Atlas.**
Representative micrographs showing the bronchial epithelia on FFPE lung tissues: immunohistochemistry for CHRNA3, CHRNA7, and CHRNB1.

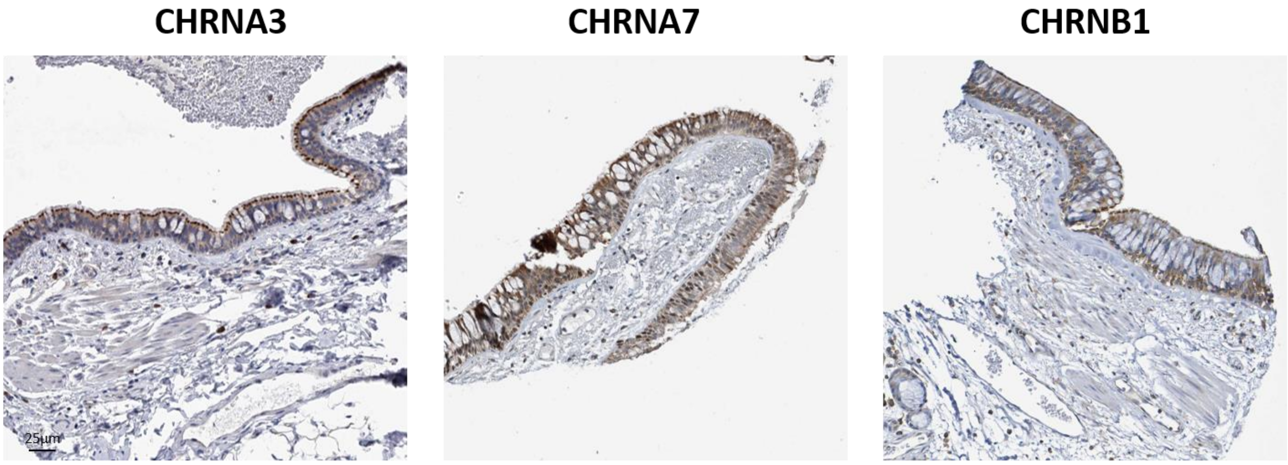
